# Genetically-encoded probes to determine nonspecific hydrophobic and electrostatic binding in cells

**DOI:** 10.1101/2023.06.27.546658

**Authors:** Weiyan Zuo, Meng-Ruo Huang, Fabian Schmitz, Arnold J. Boersma

**Affiliations:** DWI-Leibniz Institute for Interactive Materials, Forckenbeckstrasse 50, 52074 Aachen, NRW, Germany; Institute of Technical and Macromolecular Chemistry, RWTH Aachen University, Worringerweg 1, 52074 Aachen, NRW, Germany; Cellular Protein Chemistry, Bijvoet Centre for Biomolecular Research, Faculty of Science, Utrecht University, Utrecht, the Netherlands

## Abstract

Proteins interact nonspecifically with other components in the crowded cell through associative interactions. This environmental stickiness alters for example folding stability, protein diffusion, and aggregation propensity. However, the magnitude and variation in nonspecific electrostatic and hydrophobic binding energies in the cell are unclear. Here, we develop genetically-encoded fluorescence excitation ratiometric probes to determine nonspecific binding interactions. We determine hydrophobic and electrostatic interactions by systematically varying a sensing peptide on the probe. The sensors are verified in vitro and tested in HEK293T, where the nonspecific binding is highest for highly cationic and hydrophobic domains. Perturbing the cell by energy depletion increases the dependence of binding strength on peptide electrostatics, showing that the cellular conditions tune the nonspecific interaction architecture in cells. The sensors will allow estimation of nonspecific interactions and how these interactions may change in response to stresses.

## Introduction

The intracellular environment comprises proteins, polynucleotides, lipids, glycans, ions, water, and metabolites. The concentration of macromolecules is between 100-400 g/L (*1-3*). This molecular density and chemical diversity allows many nonspecific and nonfunctional molecular interactions arising from steric, electrostatics, van der Waals forces, hydrogen bonding, or hydrophobic interactions (*4*). Hence these nonspecific interactions should always be present (*5*)(*6*). The associative interaction network may be host-dependent, providing the correct environment for a native protein (*7, 8*), also known as the protein’ s quinary structure. Such interactions have thus very low specificity and dissociation constants of hundreds of micromolar to tens of millimolar (*9, 10*). In contrast, specific interactions require precise tuning of the protein interfaces to overcome the relatively low concentration of the binding partner surface area (*11, 12*).

Unwanted associative interactions, or stickiness, have major consequences as they can destabilize a protein (*13, 14*). Furthermore, misfolded proteins become sticky by exposing hydrophobic residues that recruit chaperones through attractive interactions with very low specificity. Next to protein stability (*15*) and folding (*16*), these attractive interactions also retard diffusion (*17*). Hydrophilicity and the charge determined the folding stability (*18, 19*), and probe diffusion decreases with positive charge and hydrophobicity through enhanced stickiness (*20-23*). Single point mutations can already change the stability of native proteins through interactions with the environment (*13*). Moreover, these interactions may depend on cellular conditions, as cytoskeletal depolymerization leads to increased stickiness in a protein-specific manner (*21*). Hence, nonspecific electrostatic and hydrophobic interactions are important in cells and govern protein behavior.

Nonspecific interactions can be determined with mass spectrometry combined with cross-linking or pull-down assays, either in-cell or cell lysates, gives insight into nonspecific binding (*24*). Common partners are chaperones, cytoskeletal proteins, and polynucleotide-binding proteins (25). While these measurements provide binding partners, obtaining thermodynamic data from them is challenging. In-cell thermodynamic data has been extracted from in-cell NMR of protein stability or diffusion, and fluorescence-based methods such as Förster resonance energy transfer (FRET) (*25*) to determine protein conformation, or diffusion determination through fluorescence recovery after photobleaching (FRAP) (*26*), fluorescence correlation spectroscopy (FCS) (*27*) or single-particle tracking (*28*). While these methods provided the effect of environmental stickiness on probe protein folding, diffusion, and binding, a systematic binding affinity determination as a function of hydrophobicity and electrostatics has been lacking, while being of fundamental importance.

Here we present a series of genetically-encoded fluorescent probes for detecting nonspecific interactions in vitro and in vivo. These ratiometric excitation sensors consist of short peptides with different charges and hydrophobicity linked to a circular-permuted GFP (*29*). We calibrated these sensors in different surface-modified BSA solutions to show the combined effect of hydrophobicity and electrostatics. Testing these probes in HEK293T cells reveals a nonspecific binding profile with a dependence on charge and hydrophobicity similar to wild-type BSA solutions. We show that ATP depletion with FCCP alters the interaction profile inside the cells.

## Results

To obtain a sensor for nonspecific interactions, we hypothesized that the fusion of small interacting peptides to GFP would provide a change in the excitation spectrum upon binding an analyte. Wild-type GFP has two excitation peaks at 395 nm and 488 nm that yield emission at 509 nm, which represent the different protonation states of GFP chromophore. Analyte binding to the interacting peptide could perturb the GFP structure, thereby changing the environment of the GFP chromophore and the excitation ratio (*30*). To obtain the optimal design, we used DNA-binding Lys-Trp-Lys repeat peptides (KWK)_n_ (*31, 32*). We first fused the peptide to different circular permuted GFP backbones, that is enhanced green fluorescent protein (cpEGFP), enhanced yellow fluorescent protein (cpEYFP), and green fluorescent protein (cpGFP). The backbone was split at Typ145, and the original N- and C-terminus were connected with a flexible linker (GGSGGT), similar to previous cpGFP sensors (*33, 34*). The cpEYFP backbone gave high sensitivity to DNA solutions but was also highly salt-sensitive (**Figure S2B**). In contrast, both cpEGFP and cpGFP were not sensitive to salt but also sensitive to DNA (**Figure S4A**). We selected the cpGFP because it has a more significant 405 nm excitation peak, providing a more reliable ratiometric readout because the autofluorescence in a cell is higher at this wavelength.

We made constructs encoding (KWK)_2_ peptides at both N- and C-terminus and with a (KWK)_4_ peptide at the N-terminus, respectively. The double (KWK)_2_ shows a clear ratio change with DNA concentration (**Figure S1C**). However, two interacting domains could give divalent or intramolecular interactions, and this probe was discarded. cpGFP with (KWK)_4_ displayed a clear fluorescence change when titrating DNA solutions (**Figure S1A**). The DNA binding quenches mainly the excitation at 488 nm, providing a DNA concentration-dependent increase of the 405 nm/488 nm excitation ratio (**Figure S1B**). DNA binding likely partially opens the beta-barrel and the entry of water molecules and destabilizes the chromophore state with excitation around 488 nm (*35*). As expected, the sensor also binds chemically similar molecules, such as RNA and ATP (**Figure S3**). We thus selected cpGFP fused to a 12 amino acid peptide at the N-terminus as the optimal design for nonspecific interaction sensing.

To determine binding interactions originating from charge and hydrophobicity, we substituted the sensing domain with peptides covering a range of charge and hydrophobicity (**Figure 1A**). We used lysine as a positive charge, glutamate as a negative charge, leucine as a hydrophobic residue, and serine as a hydrophilic residue. We substituted the amino acids in the repeat units, starting at GFP(KKK)_4_ and GFP(EEE)_4_ to hydrophobic GFP(KLL)_4_ or GFP(ELL)_4_ to hydrophilic GFP(SSS)_4_. Pure hydrophobic GFP(LLL)_4_ could not be overexpressed. The hydrophobicity index of the insertion peptide was calculated according to the Kyte-Doolittle scale (*36*), and the number of lysine or glutamic acid residues determined the net charge of the insertion peptide. The resulting excitation spectrum for all 11 purified sensors shows the characteristic excitation maxima at 405 and 488 nm (**Figure S5**), while the intensity of each band was sensor-dependent.

**Figure 1:**
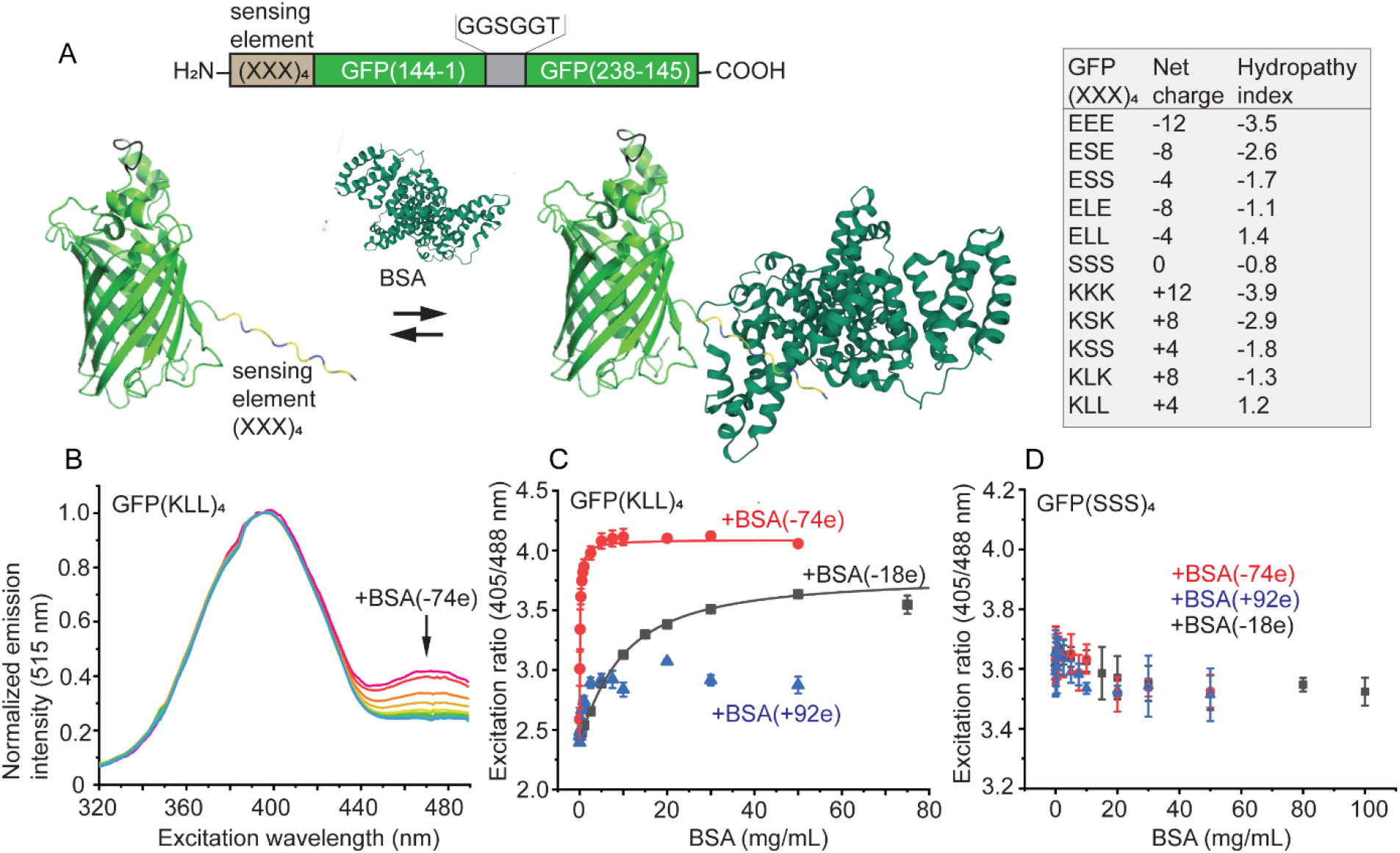
Sensing associative interactions with a ratiometric probe with a systematically varied sensing element. A. Design of the probe for nonspecific interactions and sensing concept. The sensing element domain is varied systematically to cover a range of charges and hydropathy indices. B. Normalized excitation spectrum at the emission of 509 nm for the titration of BSA(−74e) to GFP(KLL)_4_, showing the quenching of the 488 nm excitation band upon binding. C. Ratiometric readout of the titration of different modified BSAs to GFP(KLL)_4_, demonstrating the charge dependence on the binding curves. D. Titration of GFP(SSS)_4_ to the same BSAs shows that a neutral hydrophilic sensing element does not bind. All experiments were performed in triplicate. Error bars are standard deviation over the triplicate.

We first characterized the purified probes using bovine serum albumin (BSA) (**Figure 1A**). BSA is negatively charged and has a hydrophobic cleft that can bind hydrophobic peptides (*37*). Next to wild-type BSA (−18e), we applied supercharged BSAs to capture the electrostatic effect. We chemically modified 28 Lys residues on BSA with succinic acid to obtain BSA with 74 negative charges. 55 Glu/Asp residues were modified with ethylene diamine to achieve 92 positive charges. Circular dichroism shows that the main characteristics of the secondary structure of BSA are retained after these modifications, although minor changes can be observed (**Figure S8A**). Binding BSAs to the sensors changed the 488 nm band intensity, as before. We see a clear binding curve when titrating highly negative BSA(−74e) to the cationic and hydrophobic GFP(KLL)_4_. Binding to BSA(−74e) is virtually complete and much stronger than to wtBSA(−18e), as expected based on electrostatics. The BSA(+92e) shows only a very minor change in the readout at low concentration, which is likely due to impurities or misfolded BSA(+92) (**Figure 1C**). The GFP(SSS)_4_ probe, which is hydrophilic and neutral, did not sense any of the BSAs (**Figure 1D**). Therefore, the GFP(SSS)_4_ was used as a control for further in vitro and in vivo experiments. Isothermal titration calorimetry (ITC) confirms the strong binding between GFP(KLL)_4_ and BSA(−74e) and does not detect binding of the GFP(SSS)_4_ probe (**Figure S9**).

We fit the titration data with a Hill function to quantify electrostatic and hydrophobic interaction components for the binding of the peptides to BSA (**Figure S10-12**). Subsequently, we calculate the change in Gibbs free energy (ΔG) from the dissociation constant K_D_. We plotted the binding energies of the hydrophilic peptides versus BSA to determine the effect of electrostatics on BSA binding. We see that hydrophilic peptides do not bind BSA significantly, irrespective of charge, apart from the highly charged GFP(KKK)_4_ with ΔG =-17.8±3.7 kJ/mol (±s.d., n=3) (**Figure 2A,C**). In contrast to BSA(−18e), the binding to the supercharged BSAs is much more sensitive to the electrostatics. The cationic sensors have a 10-15 kJ/mol increase in binding energy with BSA(−74e), while the anionic sensors show no detectable binding. To assess the hydrophobic binding component, we compared the constructs where a leucine replaced the serine with the same net charge. (**Figure 2C**). Replacing one serine in a negatively charged background GFP(ESE)_4_ →GFP(ELE)_4_, i.e., an increase of 1.5 on the Kyte & Doolittle hydropathy scale (*36*), leads to a minute gain in binding energy with wtBSA as well as for BSA(−74e). In contrast, replacing two serines, +3.1 on the hydropathy scale, provides ∼15-20 kJ/mol for both cationic and anionic peptides for both wtBSA and BSA(−74e) (**Figure 2D**).

**Figure 2:**
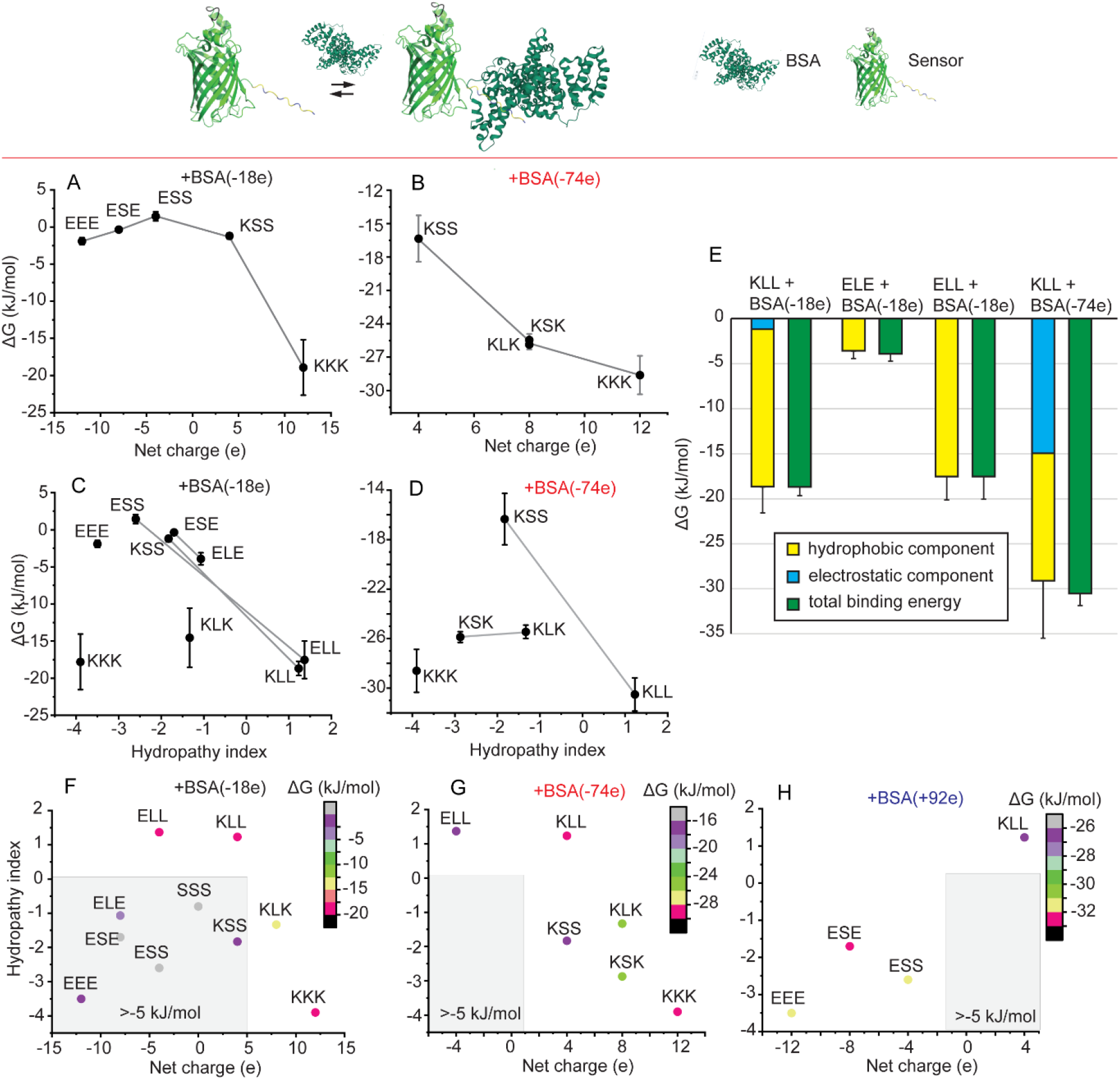
The dependence of sensor binding to BSA on charge and hydrophobicity. Dependence of the interaction energy on (A) the net peptide charge binding for hydrophilic peptides to wild-type BSA, (B) to BSA(−74e), (C) the hydropathy index on binding wild-type BSA, and (D) BSA(−74e). The lines in (A) and (B) are to guide the eye. The lines connect peptides with the same charge but different hydrophobicity in panels C and D. (E) Deconvolution of the hydrophobic and electrostatic binding energy by subtraction of the binding energy of the hydrophilic counterpart (e.g., KLL-KSS) and the opposite charge counterpart (e.g., KLL-ELL), respectively. (F-H) 2D plots on the dependence of peptide binding to both hydrophobicity and net charge versus (F) wild-type BSA, (G) negatively supercharged BSA, and (H) positively supercharged BSA. Grey shading represents the weakest binding and is to guide the eye. Grey dots are below the detection limit. All error bars are the standard deviation over three biological repeats.

To analyze whether the electrostatic and hydrophobic contributions are additive and fully account for binding, we quantified the hydrophobic component for the GFP(KLL)_4_ binding to BSA(−18e) by subtracting the binding energy of the hydrophilic counterpart, GFP(KSS)_4_ (**Figure 2E**). We assume that the difference between serine and leucine is mostly of hydrophobic origin. By subtracting the GFP(ELL)_4_ from the GFP(KLL)_4_ binding energy, we see that the electrostatics have a minor contribution and that the hydrophobic component dominates the binding. Moreover, the sum of the binding energies corresponds to the binding of GFP(KLL)_4_, suggesting that GFP(KLL)_4_ binding is the sum of the hydrophobic and electrostatic components. When executing the same procedures on the GFP(ELE)_4_ and GFP(ELL)_4_ probes, we see that the hydrophobicity fully accounts for the BSA(−18e) binding. Moreover, the hydrophobic component of GFP(ELL)_4_ and GFP(KLL)_4_ binding to BSA(−18e) are similar. As expected, when combining GFP(KLL)_4_ with BSA(−74e), the electrostatic component increases, reaching the same level as the hydrophobic component. The hydrophobic component is similar to BSA(−18e) binding, and the electrostatics and hydrophobicity fully account for the binding.

We plotted the binding energy versus the hydropathy scale and electrostatics in the same graph and see that BSA(−18e) shows a clear trend in that hydrophobic and cationic peptides bind strongly (**Figure 2F**). The amino acid that is most common in the peptides determines the high versus low binding characteristics. In contrast, the negatively charged BSA(−74e) shows already strong binding with less cationic peptides, while only the most hydrophobic of the negatively charged peptides binds, albeit much weaker than its cationic counterpart (**Figure 2G**). The same, but the inverted, trend can be seen for the BSA(+92e) (**Figure 2H**). To summarize, in all BSAs, the charge and hydrophobicity of the peptides determine the binding.

We next determined nonspecific binding inside a human cell line. We thus expressed the sensors in HEK293T cells and imaged the cells by confocal laser scanning microscopy. The sensors distributed homogeneously throughout the cell, except the hydrophobic probes GFP(KLL)_4_ and GFP(ELL)_4_ that did not partition in the nucleus, and the cationic probes GFP(KSK)_4_, GFP(KLK)_4_ and GFP(KKK)_4_ that form foci in the nuclei (**Figure 3A**). Next, we analyzed the images of each sensor and calculated the excitation ratio (405 nm/488 nm) (**Figure 3B**). To interpret the ratios in cells, we converted them to in vitro ratios using an empirically derived curve (**Figure S13**). Because water activity changes the fluorescent properties of the fluorophores to the same extent (**Figure S7**), we corrected the data with the GFP(SSS)_4_ probe, which does not bind. Moreover, the sensors were calibrated at physiological potassium chloride as this influences the charged sensors (**Figure S6A**). The sensors are not very sensitive to pH around pH 7.4 (**Figure S6B**). We find that the in-cell ratio are sensor-dependent as in buffer and we see no dependence on the sensor concentration (**Figure S16**). Therefore, the sensors do not saturate a specific binding site within this range. By calibration, we could calculate the percentage of the sensor in a bound state inside cells (**Figure 3C**). We see that highly cationic or hydrophobic sensors are 60-100% in a bound state in HEK293T cells. The other sensors are mostly 10-20% bound. Fluorescence recovery after photobleaching (FRAP) (*38*) gives recovery half times in the order of GFP(SSS)_4_ ∼ GFP(ESE)_4_ < GFP(KLL)_4_ < GFP(KKK)_4_ (**Figure S18**) showing that probe binding inversely corresponds to their diffusion as expected from increased stickiness.

**Figure 3.**
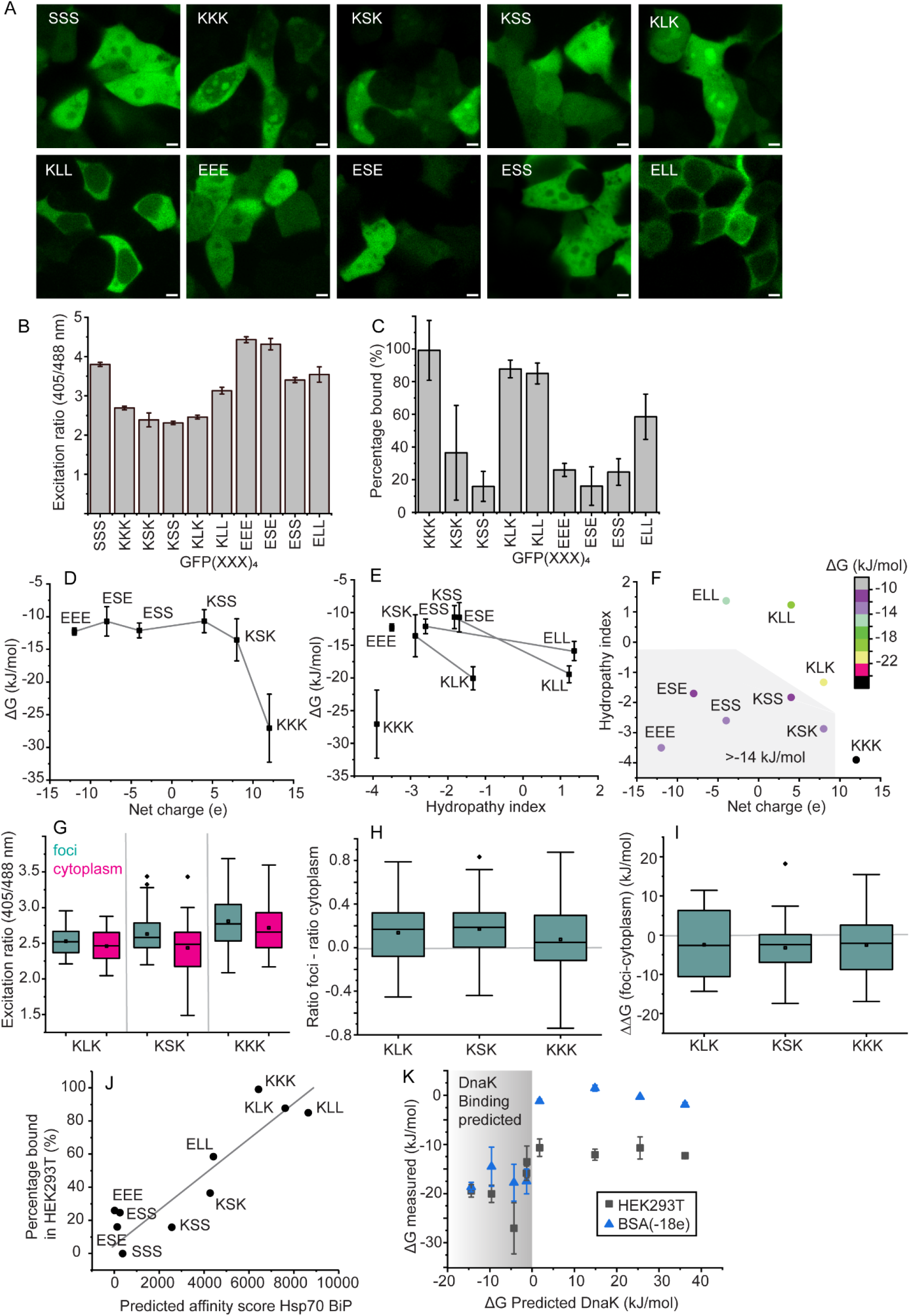
Measurement of nonspecific interactions in HEK293T. (A) Representative fluorescence confocal microscopy images of the different probes show variation in localization. White arrows indicated the locations of foci. (B) Quantification of the ratiometric readout of the probes from the cells. (C) The corresponding percentage of bound probes in the cell calculated after recalibration of the in vivo readouts. (D) Dependence of the hydrophilic peptide binding energies on their net charge. (E) Dependence of the binding energy on the hydropathy index. The lines connect peptides with the same charge but different hydrophobicity. (F) Combined plot of the dependence on charge and hydrophobicity, showing less binding for negative and hydrophilic peptides. Grey shading represents the weakest binding and is to guide the eye. (G) Comparison ratiometric readouts of all the foci compared to all the cytoplasms for foci-forming constructs. (H) Single-cell level difference in the ratio in foci versus the cytoplasm, showing a consistently higher ratio in the foci. (I) Corresponding binding energy showing a small additional stabilization of the binding in the foci. The number of cells is 40. (J) Determined binding percentages in HEK293T plotted versus predicted binding score to Hsp70 BiP in buffer (*39*). The line is to guide the eye. (K) Determined binding energy in HEK293T and in buffer with BSA(−18e) plotted versus the binding energy predicted for DnaK in buffer (*40*). The shaded area is where DnaK binding is predicted. Data points are the average over three independent replicates, and error bars are the corresponding standard deviations.

We calculated the apparent dissociation constant from the percentage of sensor bound, from which we determined the free energy of binding. To do so, we assume that the sensor binds all components in the cell equally and that the concentration of binding partners is 3 mM assuming 150 mg/mL with an average molecular mass is 50 kDa. When plotting the binding energy versus the net charge for the hydrophilic peptides, we see a curve remarkably similar to wtBSA binding (**Figure 3D**) and not the high charge sensitivity of supercharged BSAs. This suggests that the main binding partners in vivo have similar binding properties as BSA. Regarding the peptide hydrophobicity, we again see that hydrophobicity enhances binding, but the absolute effect is somewhat dampened compared to wtBSA (**Figure 3E**). The dampening effect is in absolute terms due to the charged but hydrophilic peptides displaying weak binding in HEK293T in contrast to wtBSA. Possibly, the charges are sufficient to provide at least some interactions in cells. The stronger binding peptides give roughly similar values as with wtBSA.

When combining the data in a 2D plot, we see, as expected, high qualitative similarity to wtBSA binding, suggesting that these are nonspecific binding events (**Figure 3F**). The GFP(KKK)4 shows the highest binding, GFP(KLL)_4_, GFP(KLK)_4_, GFP(ELL)_4_ intermediate, and the negative/hydrophilic constructs low binding. As chaperones are known to bind unfolded proteins that expose hydrophobic domains, we assessed if the in-cell binding could be correlated with predocted chaperone binding. To this end, we used available prediction software that estimates peptide binding to chaperones Hsp70 BiP and DnaK in buffer (*39, 40*) (**Figure 3J,K**). Indeed, we see that peptide binding in HEK293T follows Hsp70 BiP binding scores. While Hsp70 BiP itself is located in the ER, cytosolic Hsp70 could account for the binding we see. The peptides predicted to bind the bacterial DnaK, a homolog of Hsp70, also bind strongly in HEK293T and BSA(−18e). Hence, we can determine in-cell binding properties and see nonspecific patterns similar to chaperone binding. The dependence on hydrophobicity and electrostatics has high qualitative similarity to a BSA(−18e) solution.

Next, we analyzed the spatial heterogeneity in the nonspecific binding. The highly cationic constructs of two or three lysines per repeat unit form foci, possibly due to associative interactions with another biomolecule enriched in the foci, such as the polyanionic RNA in the nucleolus (*41*). We see that the excitation ratios in the foci are slightly higher than in the cytoplasm when averaging all cells (**Figure 3G**). When analyzing single cells, the ratio is, on average, ∼0.2 higher for each cell (**Figure 3H**). The higher binding affinity in the foci corresponds to roughly 2-3 kJ/mol, about 1 kBT, albeit there is significant cell-to-cell variation (**Figure 3I**). This low binding (about half a hydrogen bond per protein) is apparently sufficient to locate the probe proteins at foci.

ATP can directly interact with protein surfaces and dissipate protein self-assembly (*42, 43*). Moreover, ATP depletion and carbon starvation alter diffusion in *E. coli* (*44, 45*) and baker’ s yeast (*46*) in a probe-dependent manner (*47-49*). We thus hypothesized that ATP could direct the nonspecific interaction landscape. To test the sensors under such challenging conditions in vivo, we treated HEK293T cells with 2 μM carbonyl cyanide-p-trifluoromethoxyphenylhydrazone (FCCP) to stop ATP synthesis (*50*). We monitored the fluorescence intensity with confocal microscopy in time, and we did not observe significant changes in cell morphology for 45 minutes (**Figure 4A**). We calculated the ratio change as above (**Figure 4B**) and found that FCCP altered the ratios within 5 minutes, reaching the maximum after 5-10 minutes. The difference is not due to a temporal pH change, as the control construct GFP(SSS)_4_ did not change, indicating that the changes observed were due to the peptide domains and not the GFP. ATP does not bind the neutral or negatively charged sensors, and although it binds cationic sensors (**Figure S15**), this results in a small ratio increase with more ATP. In contrast, we see an increase in ratio upon energy depletion.

**Figure 4.**
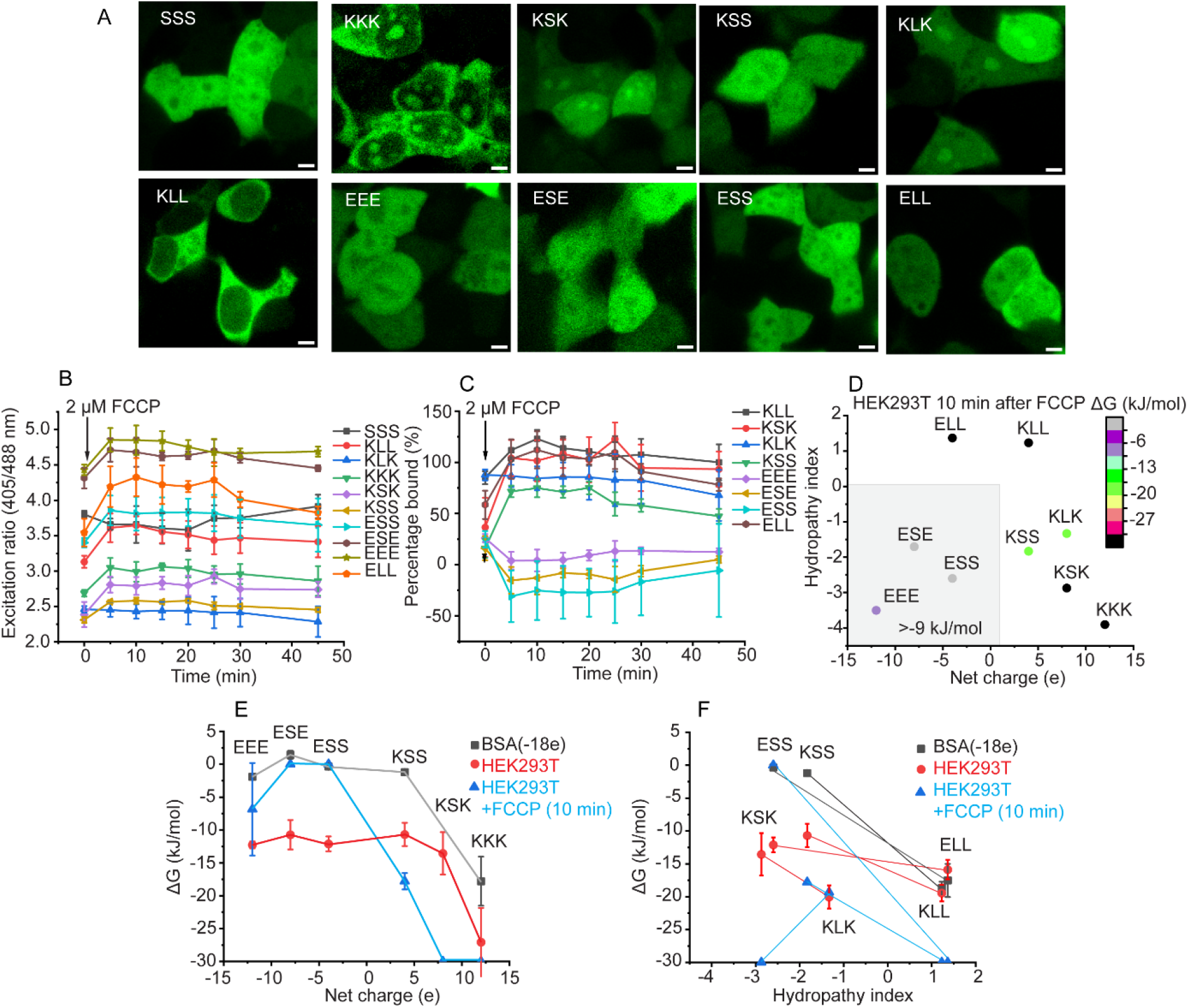
The change in the nonspecific binding profile upon FCCP treatment of HEK293T cells. A. Representative fluorescence confocal microscopy images of HEK293T cells 10 minutes after FCCP treatment. B. Corresponding ratiometric readout followed in time for the different probes. C. Calculated percentage probe bound after calibration of the in vivo data. D. Dependence of the peptide binding on the charge and hydrophobicity 10 minutes after FCCP treatment. Grey shading represents the weakest binding and is to guide the eye. Grey data points have binding affinities below the detection limit, and black data have binding affinities beyond the detection limit. E. Dependence of the peptide charge or F. of the hydrophobicity on binding to BSA, HEK293T cytoplasm, and after energy depletion. Data at -30 kJ/mol is beyond the detection limit. Lines are to guide the eye. All experiments are the average of three independent replicates, and the error bar represents the standard deviation for the replicates.

When calculating the percentage of bound probe and corresponding Gibbs free energy of binding, we see that the readout for strong-binding cationic and hydrophobic domains is saturated and the probes are fully bound. Hence, ATP depletion increases the binding energy between the cationic/hydrophobic domains and the components of the cytosol. Interestingly, the negative/hydrophilic peptides show the opposite trend, displaying lower binding potentials upon ATP removal below the detection limit (**Figure 4D**) and hence very low binding. In all cases, longer incubation leads to only minor changes in the binding potential (**Figure 4C**). When taking the cationic probes, we see that all cationic probes have higher binding, including the weakly charged GFP(KSS)4, which is more similar to the binding to BSA(−74e), and therefore the probes may bind strongly to negatively charged biomacromolecules in place of ATP. We confirmed the findings with FRAP (**Figure S18**), showing that especially the GFP(KLL)_4_ increases its recovery half time, whereas the already fully bound GFP(KKK)_4_ or the sparingly bound GFP(ESE)_4_ and GFP(SSS)_4_ showed little change. The chaperone binding predictions follow largely the experimental data, albeit with some peptides deviating (**Figure S14**). The binding differences between the peptides are enlarged due to ATP depletion, showing that the cellular conditions determine the nonspecific interaction profile of the cytosol.

## Discussion

Nonspecific and sticky hydrophobic and electrostatic interactions determine proteome stability and modulate crowding effects. Here, we developed a series of genetically-encoded fluorescent ratiometric probes to determine these nonspecific interactions in vitro and in vivo. The probes show that nonspecific interactions are most pronounced for peptides with high net positive charge or positive hydropathy index in the HEK293T cytoplasm and a simple wild-type BSA solution, with a similar dependence on peptide charge and hydrophobicity. The nonspecific interactions are modulated upon ATP depletion, where cationic peptides bind stronger and negative peptides bind weaker.

Fluorescence excitation ratiometric imaging with a single fluorophore is used frequently because it has the advantage of precise measurements and straightforward analyses. However, such sensors require controls to appreciate artifacts from pH, ions, or water activity (*29, 51*). We can correct our data with the nonbinding mutant, GFP(SSS)_4_. The resolution of the sensors is highest in the ∼5-95% bound state, corresponding to <30 kJ/mol at an estimate of 150 mg/mL (3mM) protein in the cell. The sensors need to be calibrated individually for quantification, as the response of the GFP(XXX)_4_ depends on the peptide. Likely, the sensitivity and detection range of the probes can be optimized further with the amino acids between the GFP and the sensing peptide or by applying a different fluorescent protein altogether. Nonetheless, the probes provide an interaction profile in time with minimal perturbation and with high resolution and selectivity. This is challenging to perform with other methods, if possible at all. The high binding affinity of peptides with a net positive charge has been described before, where cationic proteins and peptides have slower diffusion than their neutral counterparts, and it has been shown that protein folding stability is reduced through nonspecific or quinary hydrophobic and electrostatic interactions with the proteome (*13, 19, 52*). Usually, the binding partner is unknown, and the findings are explained by the predominantly negative charge of the biomacromolecules in cells (*53*). A lower diffusion of cationic peptides of +7e and +15e charge fused to mEos2 has been seen (*52*), where the diffusion was further slowed down in the nucleolus. Since we see that the cationic (higher than +8e) peptides locate in foci in the nucleus, this may be due to binding the polynucleotides in the nucleolus (*41*) and thereby lowering diffusion. We further see that hydrophobic peptides are excluded from the nucleus. At the same time, the nuclear pore complex should be more permeable to hydrophobic proteins (*54*): Possibly, the high binding to cytosolic components, such as chaperones, reduces the partitioning into the nucleus. Given the similarity of binding in cells to wtBSA and predictions for DnaK and Hsp70 BiP binding, the binding will be of very low specificity and probably binds the abundant “CRAPome” consisting of chaperones and cytoskeletal proteins (*55*) as well as RNA or ribosomes (*20*) in the cytoplasm. This is in line with a recently proposed general toxicity for polycationic peptides (*56*), and we show that that >80% (K_D_ = 0.5 mM) of such peptides are bound in HEK293T, depending on their hydrophobic context. Thus, the cationic and hydrophobic peptide binding is largely nonspecific and can induce peptide localization.

Besides its usual role in cells, ATP interacts directly with proteins in vitro (*43, 57, 58*), which can disperse protein assemblies. Here we show that FCCP treatment indeed alters the interaction profile in the cell. Therefore, ATP may change (nonspecific) protein-protein interactions and the organization of the cytoplasm. We see a stronger electrostatic binding component upon ATP depletion as all the cationic peptides show high binding, similar to switching from BSA (−17e) to BSA (−74e) solutions. Possibly, the dominant binding partner may also be more negatively charged such as RNA. However, other mechanisms, such as a response from the proteostasis network due to FCCP toxicity, cannot be excluded: The uncoupling of the mitochondrial membrane potential prevents the uptake of its precursor proteins that accumulate in the cytoplasm (*59*). Moreover, ATP hydrolysis usually stimulates client release from chaperones, and ATP depletion could result in irreversible complexes. Regardless of the precise mechanism, there appears to be a difference in the interaction profile of the cytoplasm due to ATP depletion, offering an explanation as to why carbon depletion in yeast retards the diffusion of some tracers and enhances the diffusion of others (*47*).

Together, our method provides a general platform to quantify nonspecific interactions in vitro and in vivo under different physiological conditions. The fluorescent proteins allow future use in specific positions inside cells by fusion with localization tags. We can obtain binding energy profiles for nonspecific interactions in vivo, giving insight into the importance of nonspecific interactions for biologics and evolution.

## Acknowledgements

The work was funded by the ERC Consolidator Grant (PArtCell; no. 864528), and the Netherlands Organization for Scientific Research (NWO Vidi; 723.015.002).

