## Supplementary figures and methods for "Genetically-encoded probes to determine nonspecific hydrophobic and electrostatic binding in cells"

### Materials and Methods

**Gene expression and purification of the non-specific interaction probes.** The synthetic gene encoding each probe was purchased from Integrated DNA Technologies (IDT, Coralville USA) and cloned into a pRSET A plasmid between restriction sites NdeI and HindIII. The *E. coli* strain BL21 (DE3) was transformed with the plasmid. The cells were grown to an $OD_{600}$ of 0.6-0.8 in LB medium (10 g/L NaCl, 10 g/L tryptone, 5 g/L yeast extract) with 1 mg/mL ampicillin at 37 °C, after which gene expression was induced overnight with 0.1 mM isopropyl-β-d-thiogalactoside (IPTG) at 25 °C. The cells were spun down, and the pellet was resuspended in buffer (10 mM sodium phosphate (NaPi), 100 mM NaCl, 0.1 mM phenylmethylsulfonyl fluoride, pH 7.4) and lysed by Multi Cycle Cell Disrupter (Constant Systems Limited, Northants UK). The cell debris was removed by centrifugation (12,000 rpm). The supernatant was supplemented with 10 mM imidazole and purified by immobilized metal affinity chromatography (His-trap FF column, Cytiva); wash/elution buffer: 20/250 mM imidazole, 50 mM NaPi, 300 mM NaCl, pH 8.0. The sensor was further purified by Superdex 200 10/300 GL size-exclusion chromatography (Cytiva) in 10 mM NaPi, pH 7.4. Purified sensor was treated with Tobacco Etch Virus (TEV) protease for 3 hours at 30 °C to remove the His-tag. Fractions that did not bind the His-trap FF column were collected. The expression and purification were analyzed by 15% SDS-PAGE, and the bands were visualized by Coomassie blue staining. Fractions containing pure protein were aliquoted and stored at −80 °C.

**Chemical modification of BSA.**

**BSA (-74e)**. To a solution of BSA (Sigma-Aldrich, >98%, lyophilized powder; 1.0 g, 15 μmol) in 0.05 M phosphate buffer (100 mL, pH = 7.5) was slowly added succinic anhydride (354 mg, 3 mmol) at 0 °C. The pH of the solution was maintained in the range of 7.5-8.0 by adding 0.5 M sodium hydroxide solution. Upon dissolution, the solution was agitated with magnetic stirring at 4 °C for 30 min. The solution was then concentrated by ultrafiltration centrifugal tubes (Vivaspin 30,000 MWCO, 20 mL) at 4 °C to give the modified BSA. The modified lysine residues were quantified by using MALDI-TOF mass spectrometry. In comparison to BSA, the succinic-BSA conjugate shows 2,800 Da molecular weight increase, which means approximately 28 lysine groups were modified.

**BSA (+92e)**. To a solution of BSA (300 mg, 4.5 μmol) and EDCI·HCl (50 mg, 0.26 mmol) in water (15 mL) was added ethylenediamine (10% aqueous, 1 mL, 1.5 mmol) at room temperature. The pH of the solution was maintained in the range of 4.5-5.0 by adding 1 N hydrochloric acid. The solution was left at room temperature for 2 h, after wich a second batch of EDCI·HCl (25 mg, 0.13 mmol) was added and the pH of the solution was readjusted to 4.5-5.0. The mixture was left at room temperature for 18 h and then concentrated by ultrafiltration centrifugal tubes (Vivaspin 30,000 MWCO, 20 mL) at 4 °C to give the modified BSA. The modified carboxylic acids were quantified by using MALDI-TOF mass spectrometry. In comparison to BSA, the ethylenediamine-BSA conjugate shows 2,517 Da molecular weight increase, which means approximately 55 carboxylic acids were modified.

**In vitro characterization of the probes.**

**DNA titration**. DNA solution (salmon sperm DNA, Invitrogen, Waltham USA) was diluted to the desired concentration with NaPi buffer (10 mM, pH 7.4). A 100 μL solution with the desired DNA concentration and purified probe was placed in a quartz cuvette and its fluorescence excitation spectrum at 520 nm emission was recorded on a Horiba Fluoromax-4P spectrometer at 25 °C. DNA fluorescence background spectrum from DNA solution without probe was recorded separately and subtracted.

**BSA titration**. Wild-type (Sigma-Aldrich, St. Louis USA) and surface-modified BSA was dissolved in NaPi buffer (10 mM NaPi, 100 mM KCl, pH 7.4) to the desired concentration. 100 μL solutions containing probe (1 μM) and BSA at the desired concentrations were placed in a 96-well plate and mixed thoroughly by pipette. The plate was scanned immediately by a SpectraMax iD3 Microplate Reader (Molecular Devices, San Jose USA). Each sample was measured in random order with equilibration up to 40 minutes. The excitation spectra at 520 nm emission were recorded at room temperature. The corresponding background spectrum from BSA solutions without probes was subtracted. The ratio ($intensity_{405nm}/intensity_{488nm}$) was plotted against BSA concentrations. Subsequently, the data was fitted to a Hill function;

$$y = ratio_{START} + (ratio_{END} - ratio_{START}) * x^n \ / \ (k^n + x^n). \qquad (S1)$$

where $K_D = k^n$ (*1*). Subsequently, the free energy of binding was calculated with

$$\Delta G = RT \ln(K_D/C^{\ominus}), \quad C^{\ominus} = 1 \text{ mol/L} \qquad (S2)$$

**Circular dichroism (CD) measurements.** The CD spectra of BSA (-18e), BSA (-74e) and BSA (+92e) were recorded using a J-1500 CD Spectrometer (JASCO, Easton USA). The protein concentrations were 0.1 mg/mL (1.5 μM). The following parameters were used: CD scale 200 mdeg/1.0 dOD, D.I.T. 2 sec, Bandwidth 1 nm. All measurements were three independent repeats.

**ITC measurements.** ITC measurements were performed on a MicroCal PEAQ-ITC (Malvern Panalytical, Malvern UK) at 25 °C. Purified probes ($GFP(KLL)_4$) and BSA (-74e) were dissolved in 10 mM NaPi, 100 mM KCl, pH = 7.4. All solutions were degassed before titrations. The sample cell was filled with 180 μL BSA (-74e) (12.9 μM), and the syringe was filled with 40 μL probe $GFP(KLL)_4$ (42.5 μM). For each titration, 0.4 μL $GFP(KLL)_4$ was first injected into the sample cell, followed by 19 injections of 2 μL $GFP(KLL)_4$ at intervals of 60 s. To subtract the heat of the dilution, the ITC buffer was injected into the protein solution under the same conditions. The raw titration data were analyzed with the MicroCal PEAQ-FITC software. The calculated $K_D$ value was the average of three independent measurements.

**Transfection of HEK293T cells**. HEK293T cells were cultured in DMEM medium containing 10% fetal bovine serum (FBS) and 1% penicillin-streptomycin (P/S) at 37 °C with 5% $CO_2$. For transfection, cells were seeded into 8-well chamber slides (Ibidi, Gräfelfing Germany), 6.7x$10^4$ cells in 200 µL growth medium (DMEM+10% FBA+1% P/S) for each well. The synthetic gene encoding each probe was purchased from IDT and cloned into pcDNA3.1 plasmid (between restriction sites NheI and BamHI). One day after plating, cells were transfected using Lipofectamine 2000 (ThermoFisher, Waltham USA) according to the manufacturer's instruction. 1 µL Lipofectamine and 250 ng pcDNA3.1 plasmid encoding the probe were diluted with 25 µL Opti-MEM I Reduced Serum Medium (Gibco) separately and incubated for 5 min at room temperature. Then, the mixture of diluted DNA and Lipofectamine (50 µL) was incubated at room temperature for 20 min and subsequently added to 150 µL DMEM (serum-free, antibiotic-free). The cell culture medium in the slide wasxchanged with the transfection medium (200 µL). The transfection medium was exchanged for complete growth medium after six hours incubation and incubated for 24h before measurement.

**Confocal fluorescence microscopy of HEK293T cells.** Before imaging, cell medium was replaced with fresh DMEM (with HEPES, no phenol red, Gibco). The cells were imaged by confocal microscope (Leica SP8, Wetzlar Germany) with a 63x/1.35 water-immersion objective at 37 °C. The probes were excited using a 405 nm LED and a 488 nm argon laser (laser power 10%) separately and fluorescence signals were recorded between 510-550 nm with a PMT detector accordingly.

For the ATP depletion experiment, carbonyl cyanide-p-trifluoromethoxyphenylhydrazone (FCCP, Sigma-Aldrich) was dissolved in DMSO (10 mM) and subsequently diluted to 2 µM with DMEM medium (with HEPES, no phenol red, Gibco). Before imaging, cell medium was replaced with DMEM (with HEPES, no phenol red, Gibco) containing 2 µM FCCP. It was previously shown that 2 µM FCCP was sufficient to deplete the cells of ATP (*2, 3*). Cells were imaged by the same method as described every 5 minutes.

The fluorescence intensity of the cells was determined in ImageJ. The intensity of blank cell cytoplasm from each image was set as background and subtracted from the measured fluorescence intensities. The 405 nm channel intensity was plotted versus 488 nm channel intensity for each cell. Linear fits with $R^2$>0.99 were obtained with the intercepts set as zero. The slope was taken as the ratio. The ratios were calibrated with the in vitro data using the empirical curve of **Supplementary Figure 13**. The percentage of probe bound inside cells was calculated using:

$$\text{Percentage bound} = |\text{ratio}_{\text{bound}} - \text{ratio}_{\text{cell}}| / |\text{ratio}_{\text{bound}} - \text{ratio}_{\text{unbound}}| \quad \text{(S3)}$$

Where $ratio_{cell}$ is the measured ratio in cells, the $ratio_{bound}$ the ratio obtained when the probe is fully bound during the BSA titration, and $ratio_{unbound}$ the ratio in of the probe pure buffer. Using the percentage of probe bound, the dissociation constant was determined as follows:

$$\text{Percentage bound} = [\text{binding partner}]_{total} / ([\text{binding partner}]_{total} + K_D) \quad (S4)$$

With $[\text{binding partner}]_{total}$ the estimated concentration of proteins in the cell, which we assumed to be 3mM. The free energy of binding inside cells was calculated according to function S2 (*4*).

**FRAP measurements.** The FRAP measurements were performed with the same microscope. The bleach area was set as 3.5 µm width, 3.37 µm height and 10.79 µm perimeter for each cell at cytoplasm and was bleached for 0.18 s with 100% LED power at 405 nm. We used the 405 nm LED to bleach, as this is the main excitation band, with the drawback of incomplete bleaching due to limited LED power. The data is highly reproducible suggesting absence of photoswitching. The data was exported from Leica software LAS X. The normalized intensity was plotted versus time and the data were fitted with one-phase exponential equation,

$$f(t)=a(1-e^{-bt})$$

The recovery half time was calculated using $\tau_{1/2} = \ln(0.5)/(-b)$. (*5*)

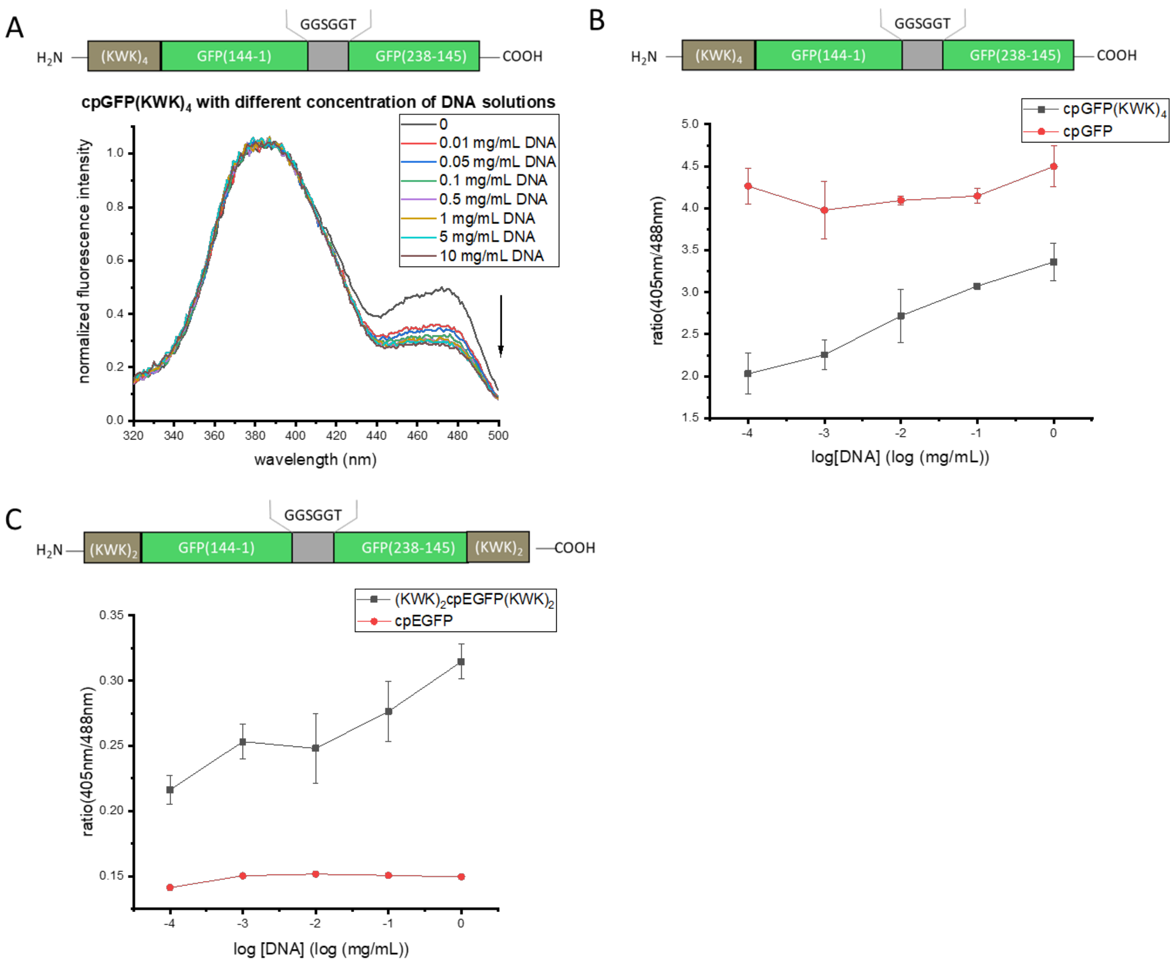


**Supplementary Figure 1:** DNA titration experiment using a Horiba Fluoromax-4P spectrometer to determine the optimal sensor design. A. Normalized excitation spectra (emission 520 nm) of cpGFP(KWK)$_4$ upon salmon testes DNA (st-DNA) titration. B. Excitation ratio change (405 nm/488 nm) of cpGFP(KWK)$_4$ upon titrating DNA and comparison with the cpGFP control. C. Excitation ratio change (405 nm/488 nm) of (KWK)$_2$cpEGFP(KWK)$_2$ upon titrating DNA and compared with cpEGFP control. Buffer is 10 mM NaPi, pH 7.4. Data are the average of three independent experiments, and error bars are the corresponding standard deviations.

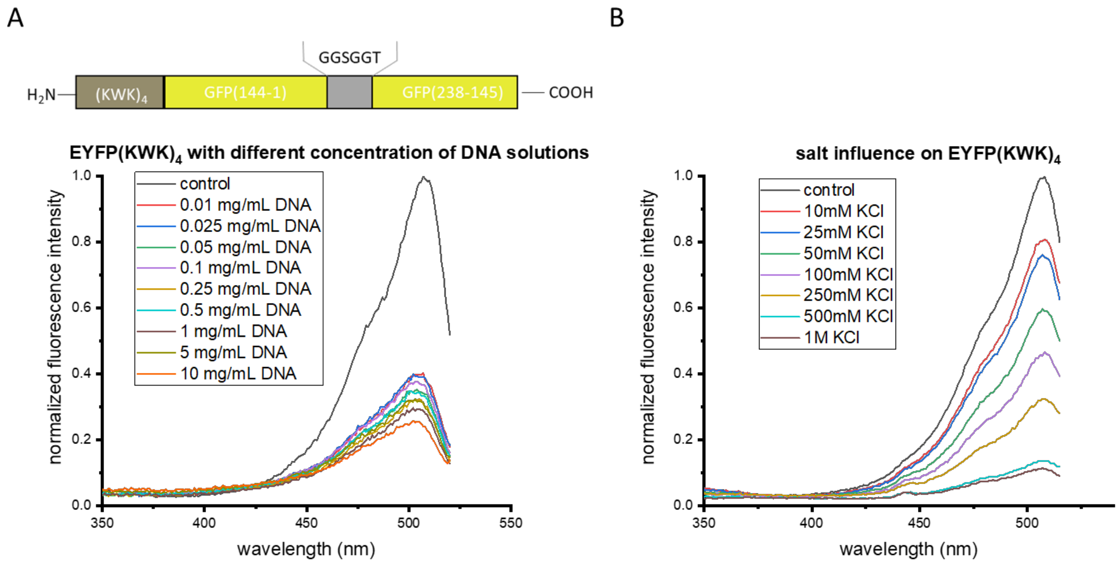


**Supplementary Figure 2**: Salt sensitivity of EYFP(KWK)4 measured by fluorescence plate reader. A. Normalized excitation spectra (emission 525 nm) of EYFP(KWK)4 upon titrating st-DNA. B. Normalized excitation spectra of EYFP(KWK)4 upon titrating KCl, showing a drastic decline in fluorescence. Buffer is 10 mM NaPi, pH 7.4.

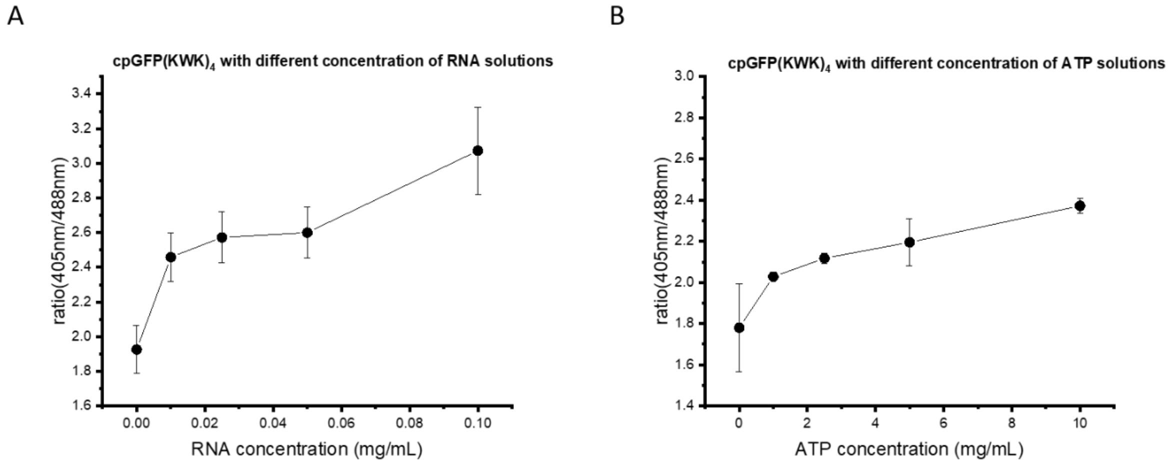


**Supplementary Figure 3**. Titration of nucleic acids to cpGFP(KWK)4 monitored by fluorescence plate reader, showing expected change in excitation ratio. A. Excitation ratio change of cpGFP(KWK)4 upon RNA (from baker's yeast, Sigma-Aldrich) titration (stock pH 7.4). B. Ratio change of sensor cpGFP(KWK)4 upon titrating ATP (stock pH 7.4). Buffer is 10 mM NaPi, pH 7.4. Data average three independent experiments, and error bars are the corresponding standard deviations.

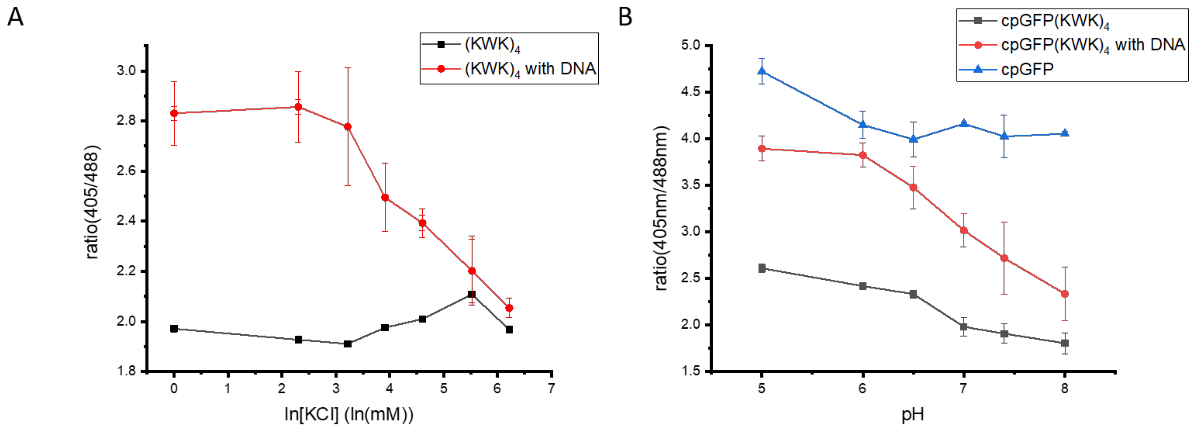


**Supplementary Figure 4**. Salt and pH influence on cpGFP(KWK)$_4$ readout. A. Excitation ratio change of cpGFP(KWK)$_4$ (black) and cpGFP(KWK)$_4$ (red) with 0.1 mg/mL DNA solution upon KCl titration. The interaction between DNA and peptide was interrupted by high concentration of KCl solution (>250 mM). B. Excitation ratio change of sensor cpGFP(KWK)$_4$ (black), sensor cpGFP(KWK)$_4$ and cpGFP control (blue) with 0.1 mg/mL DNA solution (red) in 10 mM NaPi buffer at different pH. The sensors are pH sensitive.

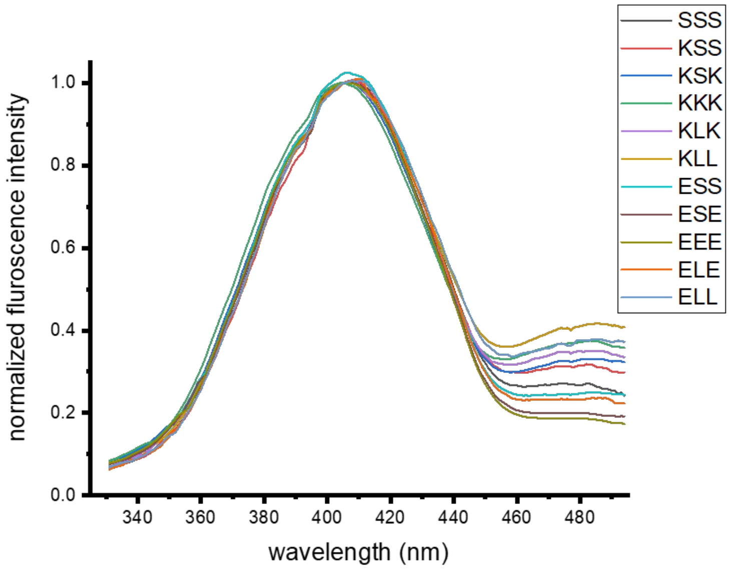


**Supplementary Figure 5**. Normalized excitation spectrum (emission 520 nm) of each sensor. All spectra were obtained in 10 mM NaPi, 100mM KCl, pH 7.4.

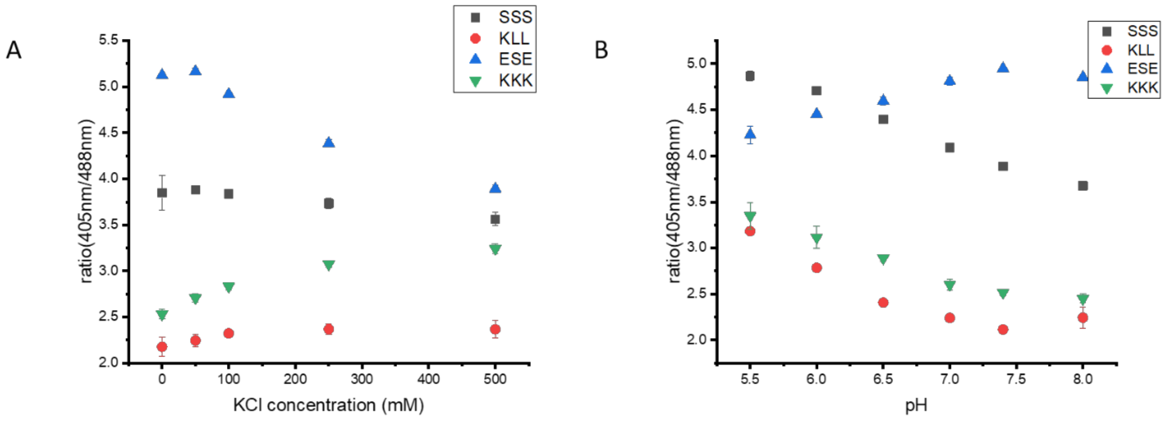


**Supplementary Figure 6**. Salt and pH influence on sensor excitation ratio change. A. Excitation ratio change of GFP(SSS)$_4$ (black), GFP(KLL)$_4$ (red), GFP(ESE)$_4$ (blue) and GFP(KKK)$_4$ (green) upon KCl titration. The sensitivity to KCl was sensor dependent. B. Excitation ratio change of GFP(SSS)$_4$ (black), GFP(KLL)$_4$ (red), GFP(ESE)$_4$ (blue) and GFP(KKK)$_4$ (green) in NaPi buffer (10 mM) at different pH. The sensors are sensitive to the pH showing that readouts need to be monitored with control sensor GFP(SSS)$_4$. Note that in-cell measurements upon ATP depletion show no change in the GFP(SSS)$_4$ probe and acidification can be excluded. All experiment were repeated as triplicates, error bars are standard deviations.

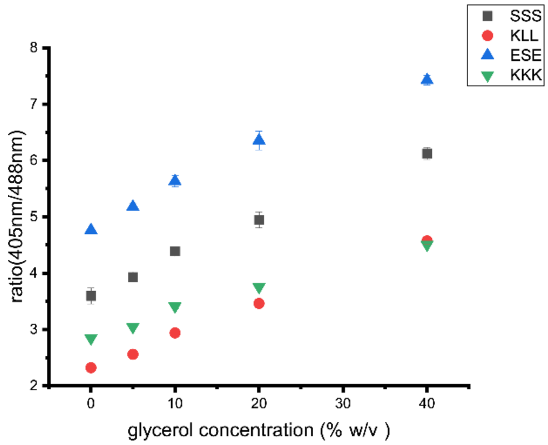


**Supplementary Figure 7**. Glycerol titration to GFP(SSS)$_4$ (black), GFP(KLL)$_4$ (red), GFP(ESE)$_4$ (blue) and GFP(KKK)$_4$ (green). Experiments in 10 mM NaPi pH 7.4. All sensors are sensitive to high glycerol content, likely from water activity, to the same extent, allowing for correcting the ratios with the GFP(SSS)$_4$ readout.

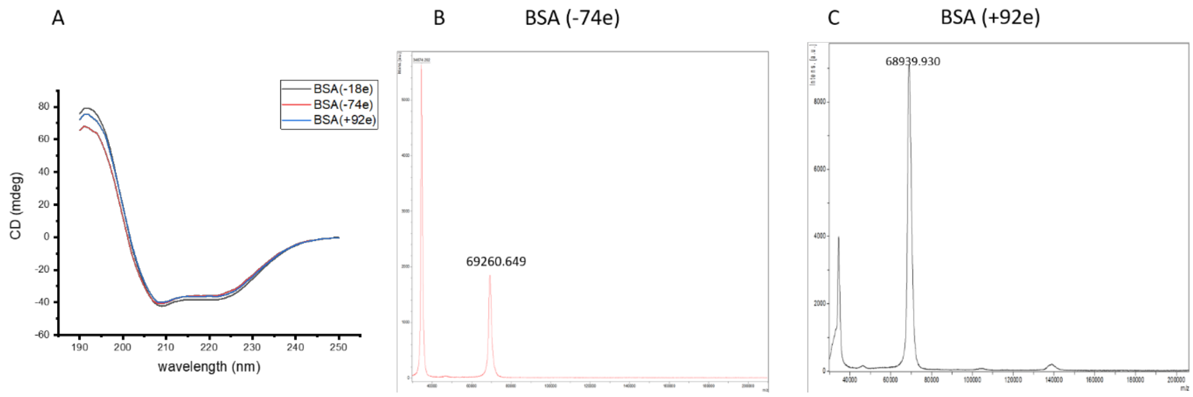


**Supplementary Figure 8**. Characterization of surface-modified BSA structure by circular dichroism and matrix-assisted laser desorption/ionization (MALDI) mass spectrometry. A. CD spectra of 0.1 mg/mL BSA in 10mM NaPi buffer at 25 °C with an HT voltage lower than 500 V. B,C. Molecular weight of BSA(-74e) and BSA(+92e) determined by MALDI-TOF mass spectrometry. Super-DHB (Merck, Germany) was selected as the matrix. MALDI-TOF mass spectrometry was performed on an ultrafleXtreme Mass spectrometer (Bruker-Daltonics, Germany).

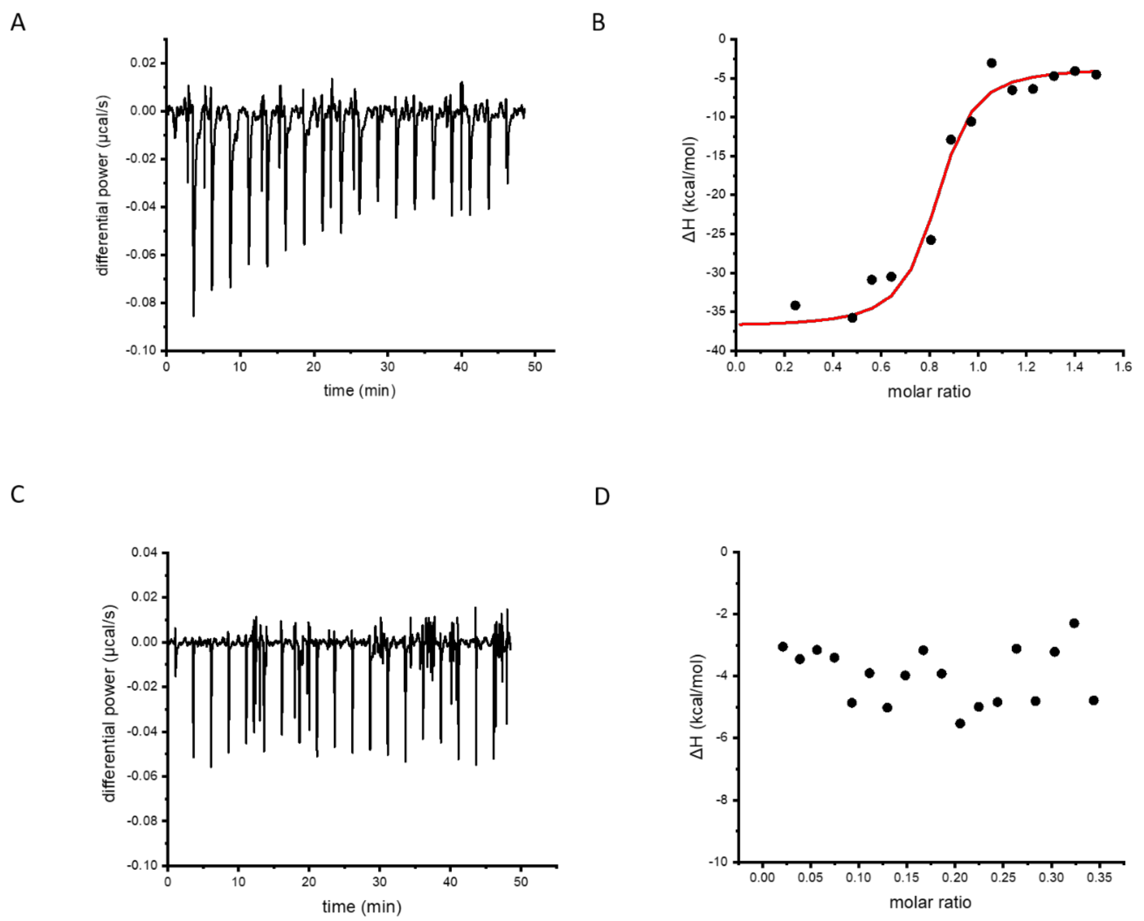


**Supplementary Figure 9**. ITC characterization of GFP$(KLL)_4$ affinity with BSA(-74e). Binding could be observed, albeit the low heat generated upon binding prevented precise $K_D$ determination. The BSA(-74e) concentration in the cell was 12.9 µM, and GFP$(KLL)_4$ concentration in the syringe was 42.5 µM. ITC characterization of GFP$(SSS)_4$ with BSA(-74e) showed no binding. All measurements were in 10 mM NaPi, 100 mM KCl, pH 7.4 and measured in triplicate.

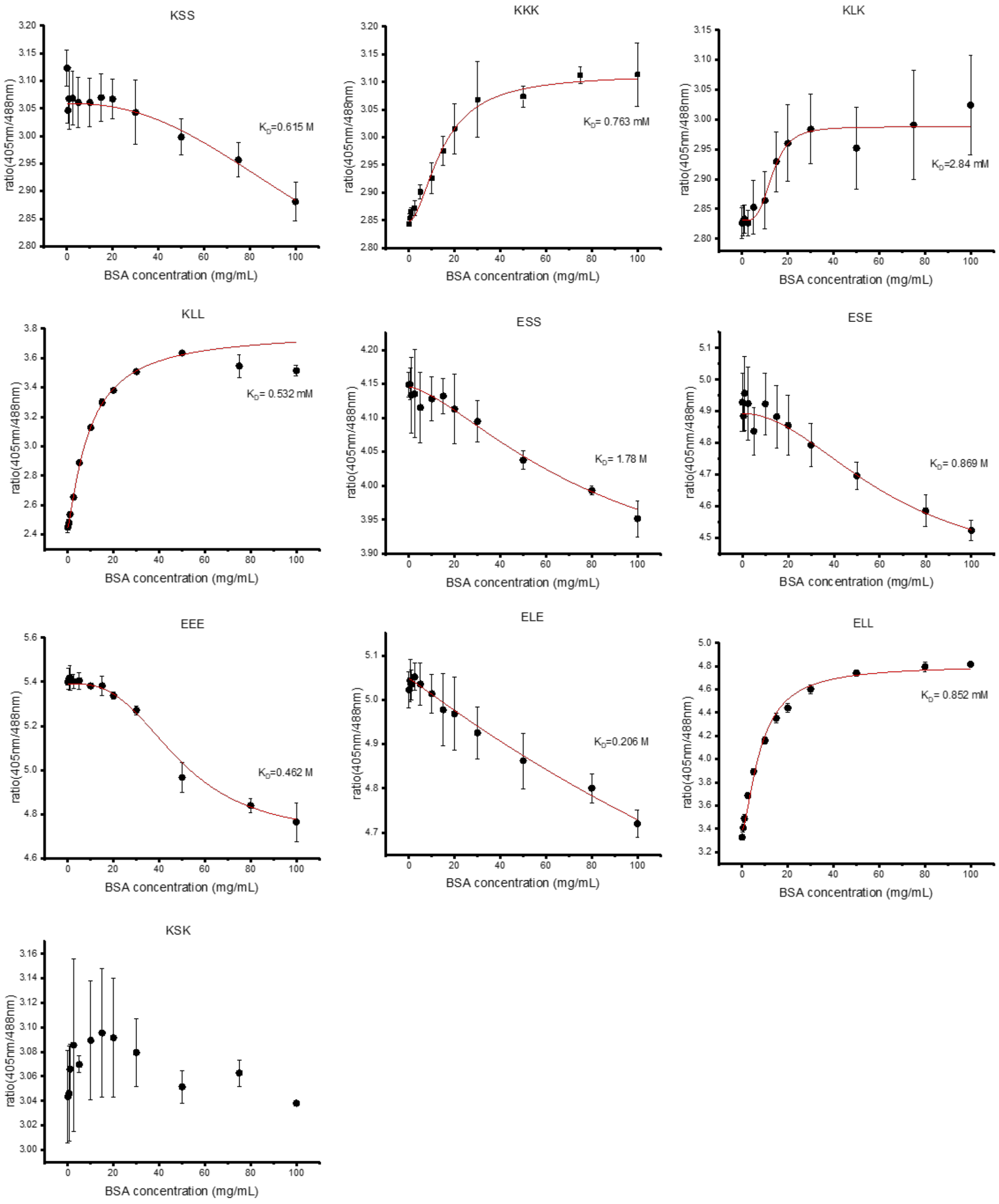


**Supplementary Figure 10**. Dose-response curves for purified sensors upon BSA (-18e) titration. All data were obtained and analyzed as described in the methods section. The excitation ratio change of GFP(KSK)$_4$ was not fit with a Hill function as the binding curve is more complex. All experiments were repeated as triplicates, and error bars are standard deviations.

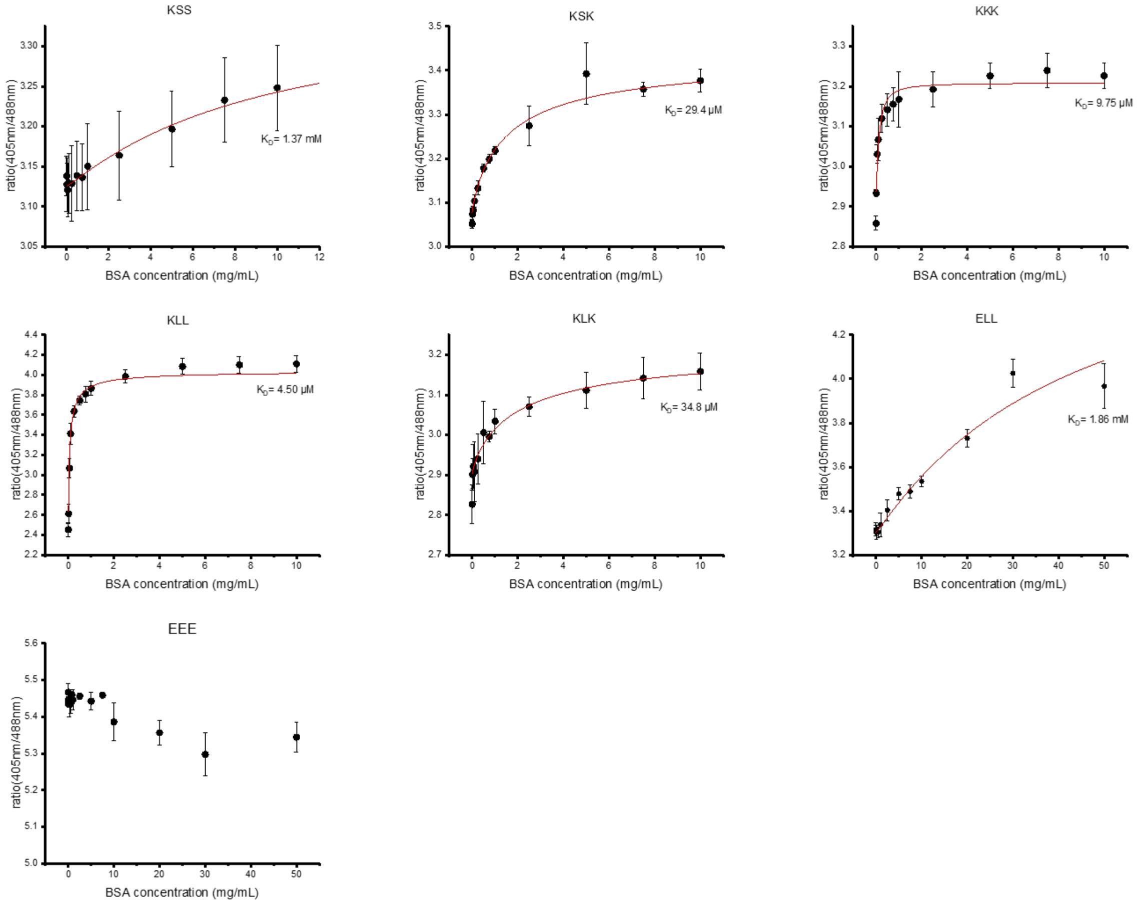


**Supplementary Figure 11**. Dose-response curves for purified sensors upon BSA (-74e) titration. All data were obtained and analyzed as described in the methods section. Excitation ratio changes of GFP(ESS)$_4$, GFP(ESE)$_4$, GFP(ELE)$_4$, and GFP(EEE)$_4$ were not fitted with a Hill function as there was no clear binding curve (data not shown for GFP(ESS)$_4$, GFP(ESE)$_4$, GFP(ELE)$_4$). All experiments were repeated as triplicates, and error bars are standard deviations.

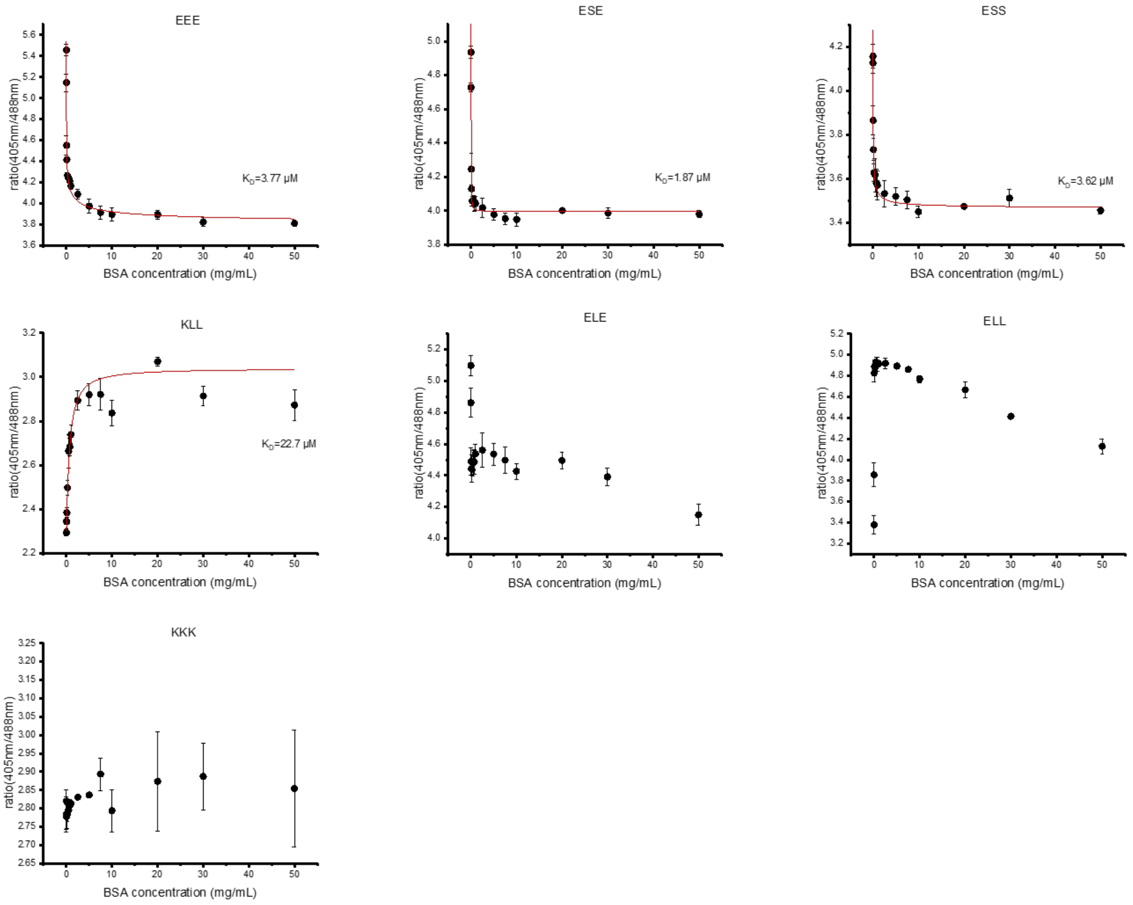


**Supplementary Figure 12**. Dose-response curves for purified sensors upon BSA (+92e) titration. All data were obtained and analyzed as described in methods section. The ratiometric changes of $GFP(KKK)_4$, $GFP(KSK)_4$, $GFP(KLK)_4$, $GFP(KSS)_4$, $GFP(ELE)_4$, $GFP(ELL)_4$ were not fitted with a Hill function due to too high binding, complex binding, or no binding (data of $GFP(KSK)_4$, $GFP(KLK)_4$, $GFP(KSS)_4$ are not shown). All experiments were repeated as triplicates, and error bars are standard deviations.

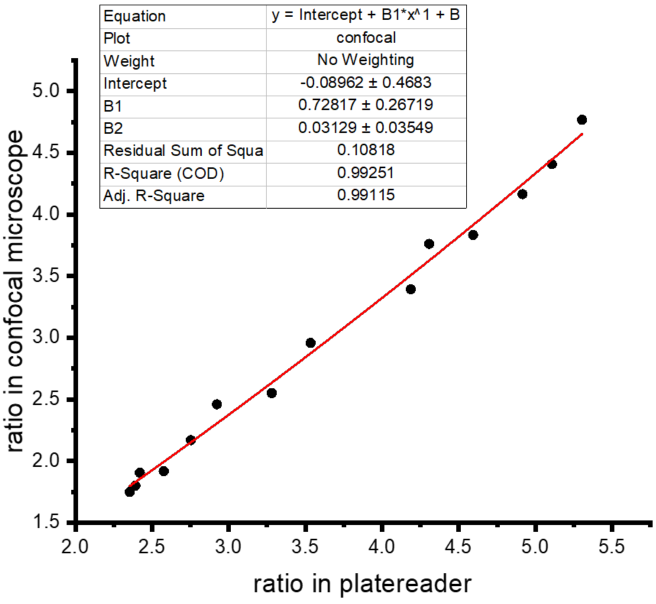


**Supplementary Figure 13**. Empirical relation between 405/488 ratios determined by fluorescence platereader and by confocal microscope.

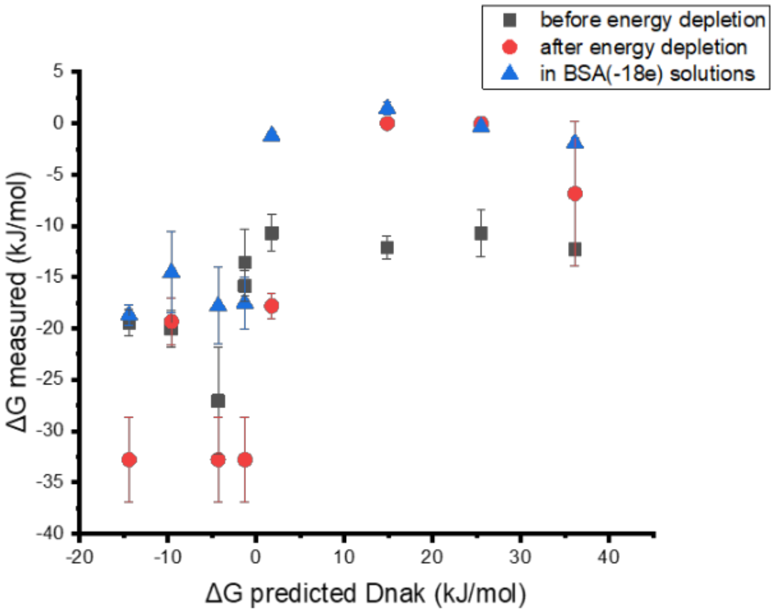


**Supplementary Figure 14**. As in Figure 3K, including ATP depletion after 10 min. FCCP.

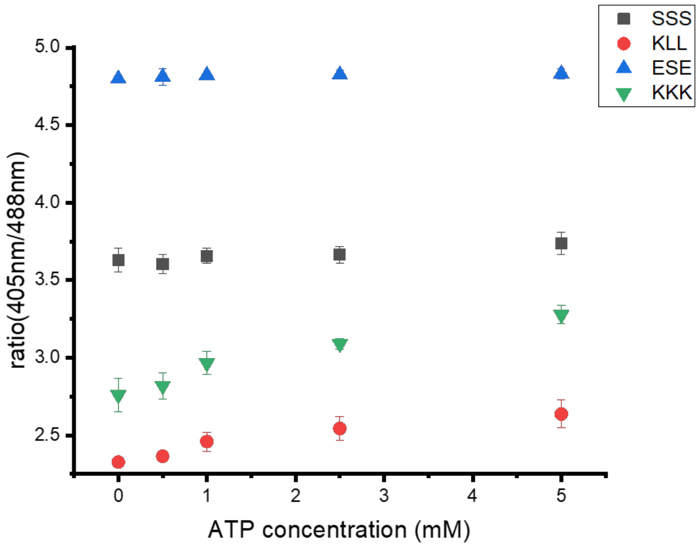


**Supplementary Figure 15**. Sensitivity of purified sensor to ATP concentration. GFP(SSS)$_4$ and GFP(ESE)$_4$ showed no excitation ratio change with ATP, while GFP(KLL)$_4$ and GFP(KKK)$_4$ showed ratio increases. Buffer is 10 mM NaPi, 100 mM KCl, pH 7.4. All experiment were repeated as triplicates, error bars are standard deviation.

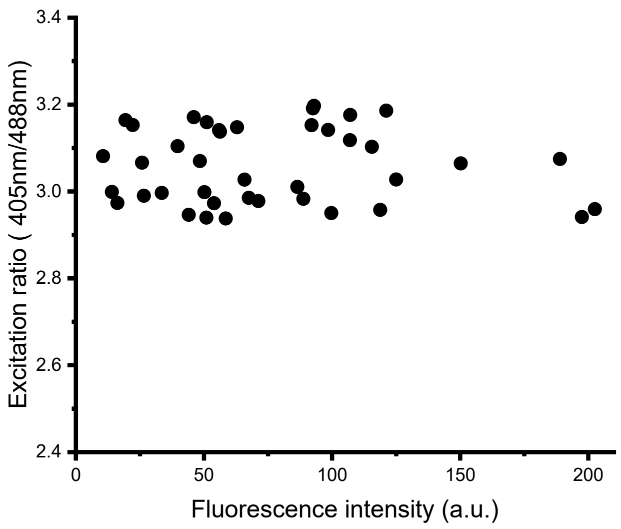


**Supplementary Figure 16.** In-cell ratio showed no dependence on sensor concentration in HEK293T cells expressing GFP(KLL)$_4$. HEK293T cells were cultured and transfected as described in the method section. Fluorescence intensities of 40 cells were recorded and analyzed as described. Excitation ratio (405nm/488nm) was plotted against the fluorescence intensity at 405 nm channel, and showed no sensor concentration (indicated by fluorescence intensity) dependence.

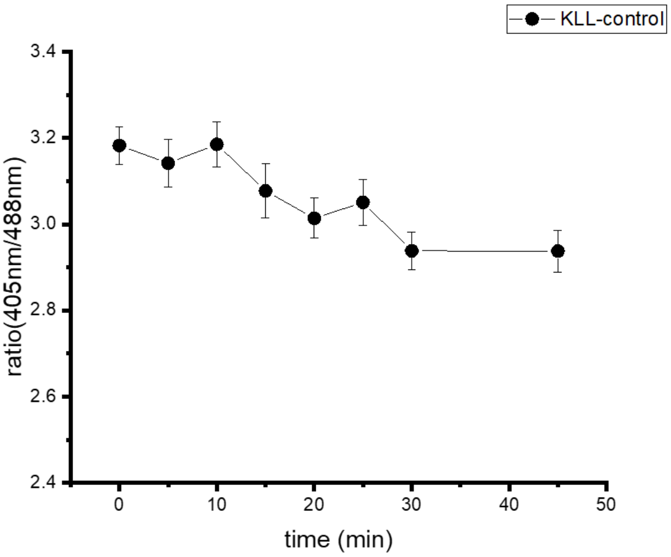


**Supplementary Figure 17**. Empty vehicle control of the addition of 0.1% (v/v) DMSO to HEK293T cells expressing GFP(KLL)$_4$. A slow and relatively small decrease in ratio was observed. HEK293T cells were cultured and transfected as described in the method section, and the culture medium was replaced with DMEM medium with 0.1% (v/v) DMSO. Errors bars represent standard deviation over three independent experiments.

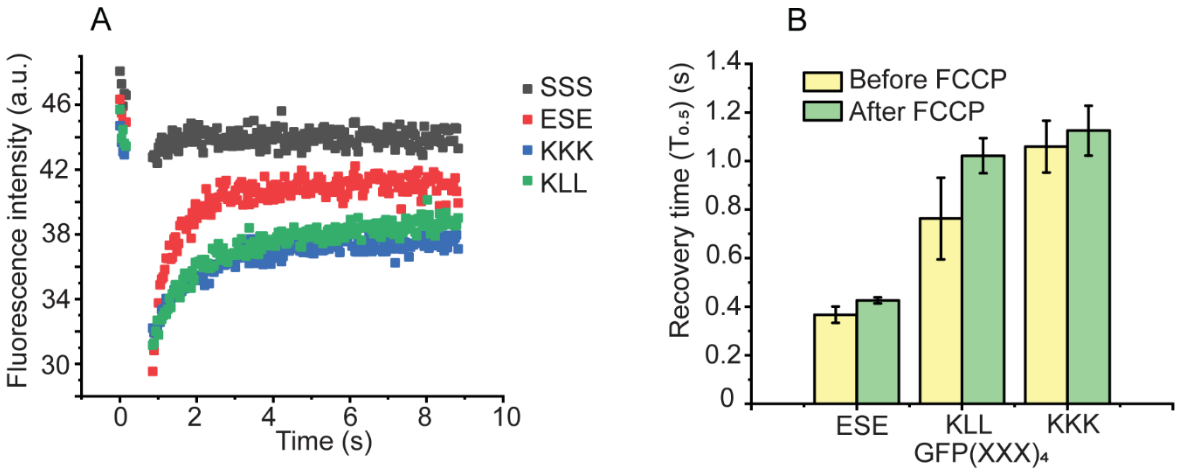


**Supplementary Figure 18**. A. Example traces of the bleaching of the probes with 405 nm LED and subsequent probe recovery. B. Quantification of the corresponding half times, showing that the stronger binding probes also diffuse slower. The GFP(SSS)$_4$ recovery is too fast for our set-up to observe the recovery curve. FRAP is after 5 minutes of FCCP treatment. All experiments are the average of three independent replicates, and the error bar represents the standard deviation for the replicates. The replicates each consisted of nine cells. The same spot was bleached three times in a all cases and showed the same recovery curve, showing that the samples do not suffer photodamage.

**Protein sequences.**

The sequences for mammalian cell expression are shown. Those for overexpression in *E. coli* contain the MHHHHHHGSGENLYFQCG sequence at N-terminus of the sequences below. In those cases the first methionine in the sequences depicted below is omitted.

**GFP(SSS)$_4$**

MSSSSSSSSSSSSAFNSHNVYITADKQKNGIKVNFTVRHNVEDGSVQLADHYQQNTPIGDGPV
LLPDNHYLSTQTVLSKDPNEKRDHMVLHEYVNAAGITHGMDELYKGGSGGTMSKGEELFTG
VVPILVELDGDVNGHKFSVRGEGEGDATNGKLTLKFICTTGKLPVPWPTLVTTLSYGVQCFSR
YPDHMKQHDFFKSAMPEGYVQERTISFKDDGTYKTRAVVKFEGDTLVNRIELKGTDFKEDG
NILGHKLEYN

**GFP(KSS)$_4$**

MKSSKSSKSSKSSAFNSHNVYITADKQKNGIKVNFTVRHNVEDGSVQLADHYQQNTPIGDGP
VLLPDNHYLSTQTVLSKDPNEKRDHMVLHEYVNAAGITHGMDELYKGGSGGTMSKGEELFT
GVVPILVELDGDVNGHKFSVRGEGEGDATNGKLTLKFICTTGKLPVPWPTLVTTLSYGVQCFS
RYPDHMKQHDFFKSAMPEGYVQERTISFKDDGTYKTRAVVKFEGDTLVNRIELKGTDFKED
GNILGHKLEYN

**GFP(KSK)$_4$**

MKSKKSKKSKKSKAFNSHNVYITADKQKNGIKVNFTVRHNVEDGSVQLADHYQQNTPIGDG
PVLLPDNHYLSTQTVLSKDPNEKRDHMVLHEYVNAAGITHGMDELYKGGSGGTMSKGEELF
TGVVPILVELDGDVNGHKFSVRGEGEGDATNGKLTLKFICTTGKLPVPWPTLVTTLSYGVQC
FSRYPDHMKQHDFFKSAMPEGYVQERTISFKDDGTYKTRAVVKFEGDTLVNRIELKGTDFKE
DGNILGHKLEYN

**GFP(KKK)$_4$**

MKKKKKKKKKKKKAFNSHNVYITADKQKNGIKVNFTVRHNVEDGSVQLADHYQQNTPIGDG
PVLLPDNHYLSTQTVLSKDPNEKRDHMVLHEYVNAAGITHGMDELYKGGSGGTMSKGEELF
TGVVPILVELDGDVNGHKFSVRGEGEGDATNGKLTLKFICTTGKLPVPWPTLVTTLSYGVQC
FSRYPDHMKQHDFFKSAMPEGYVQERTISFKDDGTYKTRAVVKFEGDTLVNRIELKGTDFKE
DGNILGHKLEYN

**GFP(KLK)$_4$**

MKLKKLKKLKKLKAFNSHNVYITADKQKNGIKVNFTVRHNVEDGSVQLADHYQQNTPIGDG
PVLLPDNHYLSTQTVLSKDPNEKRDHMVLHEYVNAAGITHGMDELYKGGSGGTMSKGEELF

TGVVPILVELDGDVNGHKFSVRGEGEGDATNGKLTLKFICTTGKLPVPWPTLVTTLSYGVQC
FSRYPDHMKQHDFFKSAMPEGYVQERTISFKDDGTYKTRAVVKFEGDTLVNRIELKGTDFKE
DGNILGHKLEYN

**GFP(KLL)$_4$**

MKLLKLLKLLKLLAFNSHNVYITADKQKNGIKVNFTVRHNVEDGSVQLADHYQQNTPIGDG
PVLLPDNHYLSTQTVLSKDPNEKRDHMVLHEYVNAAGITHGMDELYKGGSGGTMSKGEELF
TGVVPILVELDGDVNGHKFSVRGEGEGDATNGKLTLKFICTTGKLPVPWPTLVTTLSYGVQC
FSRYPDHMKQHDFFKSAMPEGYVQERTISFKDDGTYKTRAVVKFEGDTLVNRIELKGTDFKE
DGNILGHKLEYN

**GFP(ELL)$_4$**

MELLELLELLELLAFNSHNVYITADKQKNGIKVNFTVRHNVEDGSVQLADHYQQNTPIGDGP
VLLPDNHYLSTQTVLSKDPNEKRDHMVLHEYVNAAGITHGMDELYKGGSGGTMSKGEELFT
GVVPILVELDGDVNGHKFSVRGEGEGDATNGKLTLKFICTTGKLPVPWPTLVTTLSYGVQCFS
RYPDHMKQHDFFKSAMPEGYVQERTISFKDDGTYKTRAVVKFEGDTLVNRIELKGTDFKED
GNILGHKLEYN

**GFP(ELE)$_4$**

MELEELEELEELEAFNSHNVYITADKQKNGIKVNFTVRHNVEDGSVQLADHYQQNTPIGDGP
VLLPDNHYLSTQTVLSKDPNEKRDHMVLHEYVNAAGITHGMDELYKGGSGGTMSKGEELFT
GVVPILVELDGDVNGHKFSVRGEGEGDATNGKLTLKFICTTGKLPVPWPTLVTTLSYGVQCFS
RYPDHMKQHDFFKSAMPEGYVQERTISFKDDGTYKTRAVVKFEGDTLVNRIELKGTDFKED
GNILGHKLEYN

**GFP(EEE)$_4$**

MEEEEEEEEEEEEAFNSHNVYITADKQKNGIKVNFTVRHNVEDGSVQLADHYQQNTPIGDGP
VLLPDNHYLSTQTVLSKDPNEKRDHMVLHEYVNAAGITHGMDELYKGGSGGTMSKGEELFT
GVVPILVELDGDVNGHKFSVRGEGEGDATNGKLTLKFICTTGKLPVPWPTLVTTLSYGVQCFS
RYPDHMKQHDFFKSAMPEGYVQERTISFKDDGTYKTRAVVKFEGDTLVNRIELKGTDFKED
GNILGHKLEYN

**GFP(ESE)$_4$**

MESEESEESEESEAFNSHNVYITADKQKNGIKVNFTVRHNVEDGSVQLADHYQQNTPIGDGP
VLLPDNHYLSTQTVLSKDPNEKRDHMVLHEYVNAAGITHGMDELYKGGSGGTMSKGEELFT
GVVPILVELDGDVNGHKFSVRGEGEGDATNGKLTLKFICTTGKLPVPWPTLVTTLSYGVQCFS

RYPDHMKQHDFFKSAMPEGYVQERTISFKDDGTYKTRAVVKFEGDTLVNRIELKGTDFKED
GNILGHKLEYN

**GFP(ESS)$_4$**

MESSESSESSESSAFNSHNVYITADKQKNGIKVNFTVRHNVEDGSVQLADHYQQNTPIGDGPV
LLPDNHYLSTQTVLSKDPNEKRDHMVLHEYVNAAGITHGMDELYKGGSGGTMSKGEELFTG
VVPILVELDGDVNGHKFSVRGEGEGDATNGKLTLKFICTTGKLPVPWPTLVTTLSYGVQCFSR
YPDHMKQHDFFKSAMPEGYVQERTISFKDDGTYKTRAVVKFEGDTLVNRIELKGTDFKEDG
NILGHKLEYN
